# Impaired trophoblast efferocytosis by decidual macrophages in early-onset preeclampsia

**DOI:** 10.64898/2026.02.26.708195

**Authors:** Kylie BR Belchamber, Jennifer Tamblyn, Jenny Myers

## Abstract

Preeclampsia (PE) is a hypertensive disorder of pregnancy associated with inadequate trophoblast invasion, impaired spiral artery remodelling and increased trophoblast apoptosis, leading to malplacentation. Decidual macrophages are through to be key in clearing these apoptotic cells via efferocytosis, a process that normally promotes an anti-inflammatory phenotype. In PE, however, transcriptomic studies have shown that decidual macrophages exhibit a pro-inflammatory phenotype, suggesting their ability to efferocytose may be impaired. This study investigated whether PE alters *ex vivo* decidual macrophage function in placentas obtained from healthy pregnancies (n=11) or with early-onset PE (n=9). Macrophages isolated from both the decidua basalis and decidua parietalis of PE placentas demonstrated significantly reduced efferocytosis of apoptotic trophoblasts compared to healthy controls, accompanied by increased release of CXCL-8 and IL-6. Similarly, phagocytosis of *Streptococcus agalactiae* was significantly impaired by both macrophage subtypes. Analysis of macrophage scavenger receptors revealed that efferocytic macrophages upregulate receptors for ‘eat me’ signals, whereas non-efferocytic macrophages fail to do so. These findings demonstrate that decidual macrophages in the preeclamptic placenta exhibit impaired efferocytosis and phagocytosis, which may contribute to the accumulation of apoptotic trophoblasts and increased pro-inflammatory signaling. Further analysis of this defective function is vital to identify novel immunomodulatory treatments for PE.

**Graphical abstract:** In healthy pregnancy, decidual macrophage efferocytosis of apoptotic trophoblasts prevents inflammation through release of anti-inflammatory IL-10. In early-onset preeclampsia, failure to efferocytose causes a build up of apoptotic trophoblasts, and promotes inflammation through release of pro-inflammatory CXCL-8 and IL-6. Targeting decidual macrophage function may be beneficial in preeclampsia.

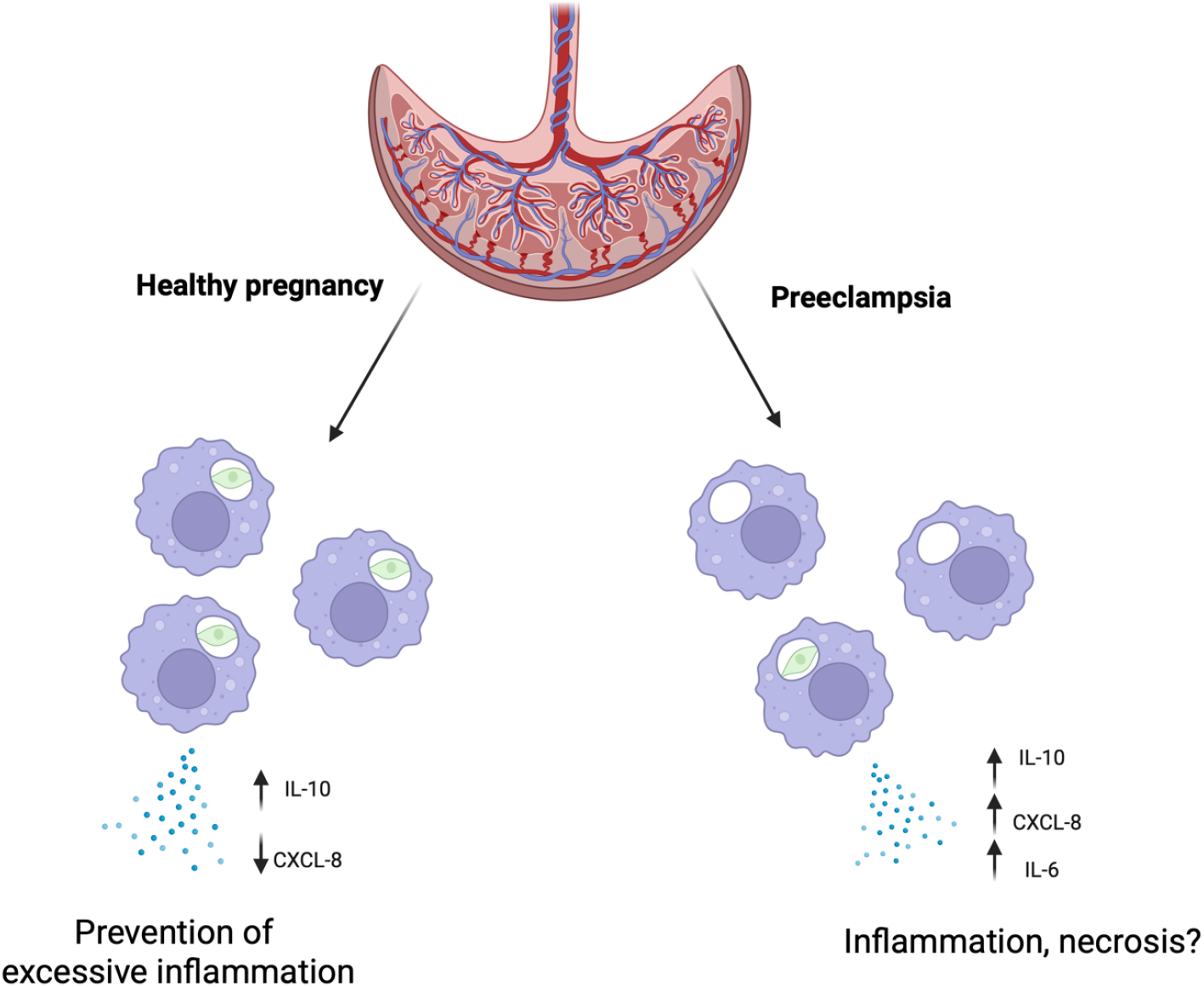

## Introduction

Efferocytosis – the process by which macrophages clear apoptotic cells - is essential for the maintenance and restoration of tissue homeostasis. Defects in efferocytosis are increasingly linked with disease pathologies, especially in tissues with high cellular turnover [1]. The placenta, which supports development of the endo-myometrium, has extremely high cellular turnover due to its rapid growth and invasion into the uterine wall, and extensive remodelling that occurs throughout pregnancy [2]. The placenta also represents a unique immunological challenge, whereby tolerance must be established to the semi-allogenic fetus, whilst dynamic inflammatory states are necessary for key stages of placental development and function. Failures in these tightly regulated processes can lead to pregnancy complications including miscarriage (up to 25% recognised pregnancies [3]), preeclampsia (affecting 5% of pregnancies [4]), fetal growth restriction (3-9% pregnancies [5]), and still birth (0.4% pregnancies [6]).

Decidual macrophages (dMφ), which originate from maternal endometrium and/or blood monocytes, are the main efferocytic cells of the maternal decidua, and appear vital in establishing materno-fetal tolerance and maintaining a homeostatic environment during pregnancy [7]. In early pregnancy, dMφ are located near the spiral arteries during trophoblast invasion and spiral artery remodelling [8], where they present an M2-like, anti-inflammatory phenotype, expressing high levels of CD206, CD163 and CD209, likely due to the abundance of M-CSF and IL-10 in the decidua [9, 10]. dMφ are also found in the decidua parietalis, where in the first trimester they express a pro-inflammatory M1-like phenotype; are able to release higher levels of CXCL-8 and 6 then decidua basalis dMφ; and are able to induce CD4+ and CD8+ T cell proliferation at a higher level than decidua basalis dMφ [11].

As pregnancy progresses, dMφ are likely essential for ongoing efferocytosis of apoptotic structural cells generated during placental growth and development [12].In the third trimester, dMφ retain an anti-inflammatory phenotype, but increase expression of phagocytic markers CD206, CD80 and HLA-DR [13]. Whilst key role in partuition are anticipated [14], functional assessment of dMφ function in pregnancy remains limited, with our understanding of their complex roles in pregnancy also restricted.

Preeclampsia (PE), a major cause of maternal and fetal morbidity and mortality, complicates >5% of pregnancies. PE is associated with defective trophoblast invasion and spiral artery remodelling, causes impaired placental vascularisation and generation of an oxidative environment [15]. Increased dMφ numbers have been reported in myometrial segments of uteroplacental arteries from PE patients compared to healthy placentas at term [16], and also within and around the spiral arteries, separating them from trophoblasts. This resembles a barrier between the invading trophoblast and spiral arteries, which may prevent the transformation of spiral arteries and lead to malplacentation [17]. Whether dMφ numbers overall change in PE is unclear, however there appears to be a pro-inflammatory environment in the PE placenta, with increased macrophage chemotactic factors CXCL-8 and MCP-1 [18-20], as well as elevated TNFα, IFNγ, IL-6 and decreased anti-inflammatory cytokine IL-10 [21]. TNFα and IFNγ induce apoptosis of extra villous trophoblasts, which may signal increased dMφ efferocytosis. However, the pro-inflammatory environment of the PE placenta may inhibit efferocytosis, leading to accumulation of necrotic cells, and promoting inflammation across gestation.

dMφ efferocytosis in the placenta is therefore likely to be vital for maintaining tissue integrity by preventing secondary necrosis, and potentially for sustaining tolerance during pregnancy. Changes in this process may be implicated in pregnancy complications. In this study, we developed a novel dMφ efferocytosis assay and sought to assess dMφ efferocytosis in healthy pregnancies, compared to those complicated by early onset preeclampsia.

## Methods

### Patient recruitment

Pregnant women were recruited to this study after providing written informed consent either from Birmingham Womens Hospital, (West Midlands ethics committee, 14/WM/1146) and St Marys Hospital Manchester, (North West ethics committee, 15/NW/0829). Healthy control participants had a singleton pregnancy with no complications as per clinician assessment, no history of PE or other pregnancy complications, and underwent caesarean section due to maternal request. Participants with PE were diagnosed between 24-32 weeks gestation by clinician according to NICE guidelines [22], including systolic blood pressure of 160mmHg or higher and proteinuria. PE participants also had a singleton pregnancy, no evidence of chronic hypertension prior to pregnancy, and no additional pregnancy complications. Participants were delivered by pre-labour caesarean section, and placentas were collected within 30 minutes of delivery and processed immediately.

### Tissue processing

Placentas were immediately processed upon receipt. To obtain decidua parietalis, the chorionic sac was scrapped using a scalpel and transferred into a glass bottle containing PBS. To obtain decidua basalis, the first 0.5cm of the maternal side of the placenta was macroscopically dissected and transferred to a second glass bottle containing PBS. These tissues were passed through a 30μm smart strainer (Miltenyi Biotech) to remove excess blood and transferred to a C-tube (Miltenyi Biotech). Samples were mechanically dissociated using a Miltenyi gentleMACS dissociator for 30 seconds. Samples were then digested using 1mg/ml collagenase V (Sigma-Aldrich) and 10mg/ml DNAse 1 (Sigma-Aldrich) for 60 minutes at 37 degrees, with agitation. The digested cell suspensions were passed through a 100μm smart strainer (Miltenyi Biotech) and washed through with PBS. Cells were centrifuged at 500g for 10 minutes to form a pellet, and resuspended in PBS. Cells were layered over a 20ml Lymphoprep (Stemcell) and centrifuged for 20 minutes at 800rpm without break. The leukocyte layer was collected and washed in PBS by centrifugation. CD14+ macrophages were isolated using the Easysep CD14+ selection kit II (Stemcell) according to manufacturer’s instructions. CD14+ cells were washed in PBS and labelled as decidua basalis macrophages (bMφ) or decidua parietals macrophages (pMφ).

### Macrophage phenotyping by flow cytometry

Freshly isolated CD14+ dMφ were fixed in 4% paraformaldehyde for 10 minutes at room temperature, then washed in PBS. Cells were resuspended in FACS buffer (PBS+0.5% BSA (Sigma Aldrich)) and blocked with 5% human serum (Sigma Aldrich) for 10 minutes, then washed in FACS buffer. Cells were incubated with antibodies (listed in table S1) for 30 minutes at 4 degrees. Cells were washed a resuspending in FACS buffer and stored at 4 degrees until analysis. Flow cytometry was performed using a BD Fortessa X20 (BD Biosciences) and analysed using Flowjo (Treestar).

### Decidual macrophage culture

bMφ or pMφ were counted using a haemocytometer and plated in 48 well plates (Corning) at 200,000 cells/well in RpMI+ media (RpMI 1640 media (Thermofischer)+ 10% FCS (Thermofischer) + 100U/ml penicillin (Thermofischer) + 100μg/ml streptomycin (Thermofischer) + 2mM L-glutamine (Thermofischer)) overnight to allow adherence. The next day, cells were gently washed and fresh RpMI+ added. Cells were then used for experiments as stated.

### Efferocytosis assay

Bewo cells (CCL-98, NTCC) were maintained in culture in T75 flasks and dMEM-F12 Hanks media (Thermofischer) + 10% FBS + 100U/ml penicillin + 100μg/ml streptomycin + essential amino acids (Thermofischer) and grown to confluence. Bewo were removed from T75 using trypsin-EDTA 0.25% solution (Thermofischer) for 5 minutes at 37 degrees, followed by quenching with dMEM-F12 media. Bewo were centrifuged at 300g for 5 minutes, and resuspended in PBS. Bewo were counted using a haemocytometer and transferred to a 6 well plate. 1ul cell tracker green CMFDA (or cell tracker DEEP red for microscopy, Thermofischer) was added, and the plate incubated under a UV lamp for 30 minutes to induce apoptosis. Apoptosis was confirmed by Annexin V/Propidium iodide staining (Supp fig 1) to ensure >80% cells were late apoptotic. Bewo were washed in PBS and resuspended in RpMI+. Media was removed from dMφ wells, and bewo added at a ratio of 4:1 (Bewo: dMφ). Plates were incubated at 37 degrees for 90 minutes, after which supernatants were removed and stored at -80 degrees for future analysis. Cells were washed gently in PBS and removed from the plate using cell dissociation buffer (Thermofischer) and vigorous pipetting. Cells were transferred to FACS tubes and fixed in 4% PFA for 10 minutes. Cells were washed and resuspended in FACS buffer for antibody staining.

### Phagocytosis assay

*Streptococcus agalactiae* (NCTC 8181) was grown overnight in 25ml Brain Heart Infusion (BHI) broth (Sigma), followed by subculture in 500ml BHI for 8 hours. Bacteria were centrifuged at 4000g for 30 minutes, and a sample taken for CFU counting on Colombia blood agar. Bacteria were heat killed at 70 degrees for 2 hours, then centrifuged, and resuspended in NaOH, and 1mg/ml Alexafluor 488 NHS Ester (Thermofischer) added. Bacteria were incubated overnight, in the dark, on a windmill rotator to allow staining. The next day, bacteria were centrifuged and resuspended in PBS. Bacteria were aliquoted to 1x10^9^CFU/ml and stored at -80. For phagocytosis assay, an aliquot of bacteria was defrosted, and sonicated for 2 minutes. Bacteria were added to dMφ at a ratio of 80:1 for 4 hours, then processed as above.

### Antibody staining

dMφ were incubated in 5% human serum for 10 minutes, then washed in FACS buffer. Cells were incubated with antibodies (listed in table S2) for 30 minutes at 4 degrees. Cells were washed a resuspending in FACS buffer and stored at 4 degrees until analysis. Flow cytometry was performed using a BD Fortessa X20 (BD Biosciences) and analysed using Flowjo (Treestar).

### Confocal microscopy

dMφ were plated on 8 well μ-Slides (Ibidi) at 100,000 cells per well, in RpMI+ and cultured overnight to allow adherence. The next day, cells were gently washed and fresh RpMI+ added. Efferocytosis or phagocytosis assays were performed as stated, then cells fixed in 10% PFA for 10 minutes. PFA was removed and cells were washed in PBS. Cells were stained with cell tracker green CMFDA for 30 minutes, and DAPI (Sigma) for 5 minutes. Cells were washed and stored in PBS until microscopy. Microscopy was performed on a Leica SP8 inverted confocal microscope and analysed using FIJI software.

### Cytokine analysis

Cytokine release was measured in supernatants using R and D duoset ELISAs for CXCL-8, IL-6, TNFα and IL-10, as per manufacturer’s instructions.

### Statistical analysis

All data was analysed using Graphpad prism software v10. For comparison of two parameters, paired or unpaired t-tests were used where appropriate. For comparison of more than two variables, a 2 way ANOVA was used. Data are presented as either individual data points plus mean, or as min-max box plots, with median.

## Results

### Patient demographics

Table 1 shows patient characteristics and pregnancy outcomes. The mothers were age matched (32-34 years), and both groups had a mix of ethnicities, and had similar gravidity and parity. PE pregnancies were delivered at an earlier gestation (average 34 weeks, n=9) compared to healthy control pregnancies (average 39 weeks gestation, p<0.001, n=11), and neonates had smaller weights (2103g vs. 3535g, p<0.001).. There was an even split of neonate sex in both groups (HC 6:5, PE 4:5 male:female).

**Table 1.**
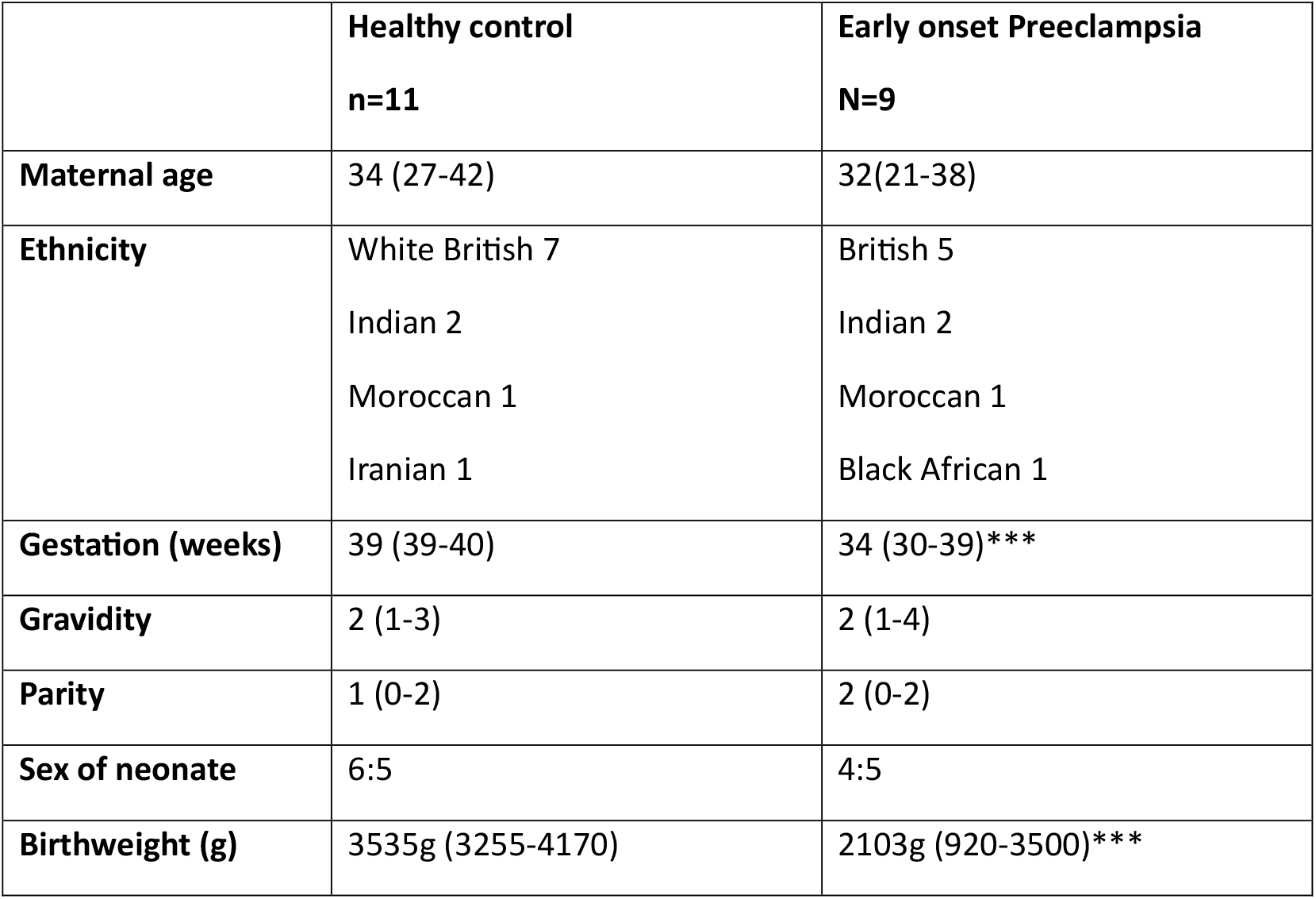
Patient demographics(median/range).

### Decidual macrophage phenotypes differ according to location

By confocal microscopy, bMφ appear rounded in shape, with multiple dendritic extensions – a typical macrophage appearance (Fig 1A). pMφ appeared morphologically different, with a more elongated shape, with long extensions (Fig 1B). This is suggestive of different phenotypes and function of these cells.

**Figure 1.**
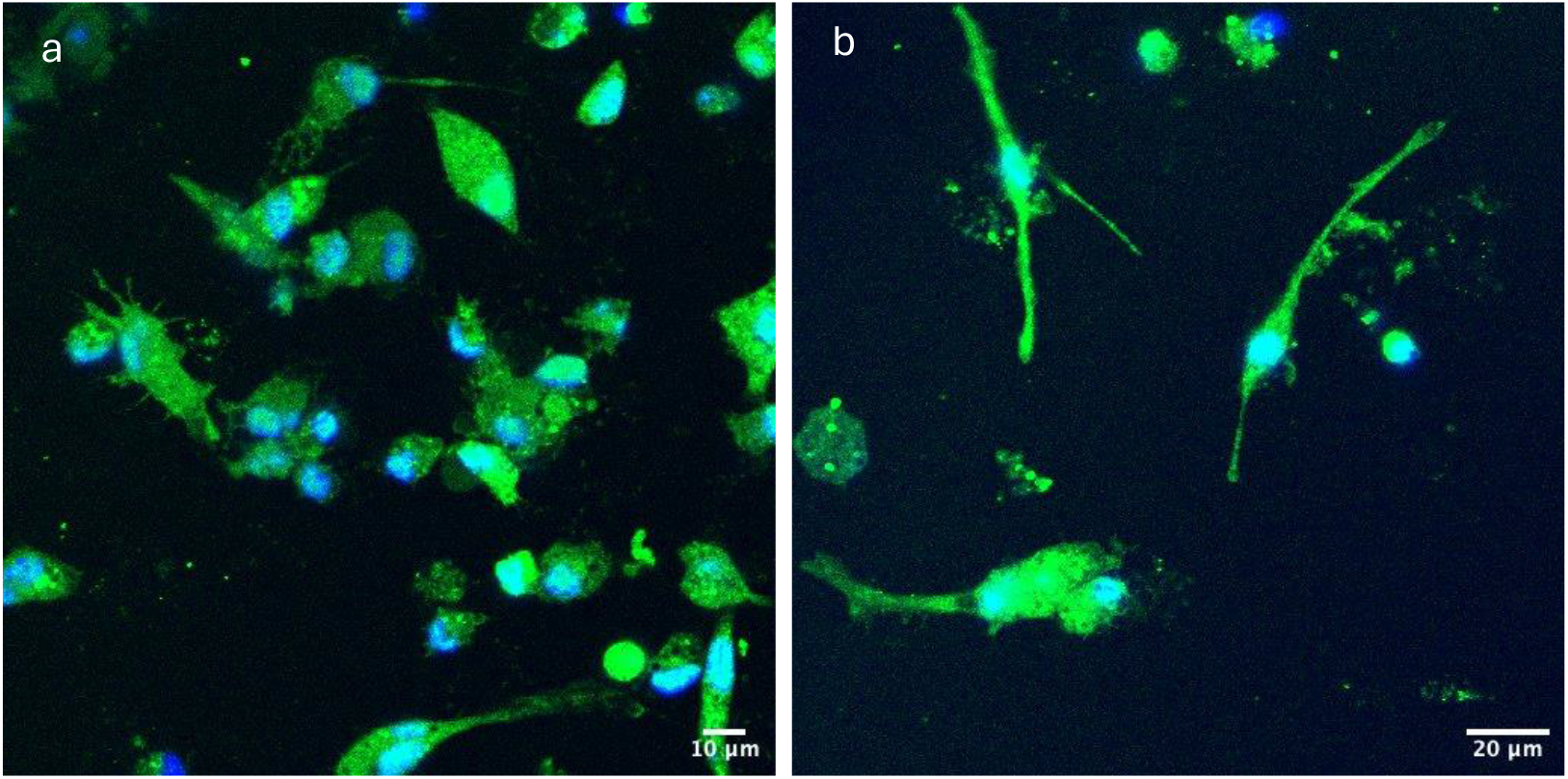
DMφ morphology. Isolated bMφ (a), and pMφ (b) stained with cell tracker green, and DAPI (nucleus) imaged by confocal microscopy.

We next defined the phenotype of dMφ from healthy pregnancies based on their location in the decidua (Table 2). Both bMφ and pMφ represented a range of phenotypes, with the majority expressing TLR4 (87% and 86%), and CD163 (79% and 81%), and lower representation of CD36+ (46% and 38%), CD206+ (40% and 53%), TLR2+ (41% and 47%) and the trophoblast receptor LLBR1+ (22% and 32%). pMφ has significantly higher levels of CD68+ cells (28% vs. 40%, p<0.01) and CD80+ cells (37% vs. 51%, p<0.05) which may indicate increased levels of pro-inflammatory skewed macrophages. The level of expression of these receptors (measured by MFI, Table 3) backed this theory up, with pMφ expressing significantly higher levels of CD68 (3.2 vs. 5, p<0.05), CD80 (3.7 vs. 5.4, p<0.05), CD206 (1.9 vs 2.9, p<0.05) and TLR4 (1.5 vs. 2.1, p<0.05). This suggests pMφ may represent a more pro-inflammatory function, primed for responses to infection, whereas bMφ are more anti-inflammatory.

**Table 2.** % receptor expression by macrophages.∗p<0.05, ∗∗p<0.05, analysed by paired t-test.

|  | BASALIS | PARIETALIS |
| --- | --- | --- |
| CD36 | 46±21 | 38±21 |
| CD68 | 28±15 | 40±23** |
| CD80 | 37±12 | 51±22* |
| CD163 | 79±25 | 81±27 |
| CD206 | 40±21 | 53±24 |
| TLR2 | 41±23 | 47±18 |
| TLR4 | 87±14 | 86±23 |
| LLBR1 | 22±19 | 32±28 |

**Table 3.** Median Fluorescent Intensity of receptor expression by macrophages. ∗p<0.05 analysed by paired t-test.

|  | BASALIS | PARIETALIS |
| --- | --- | --- |
| CD36 | 10.6±6 | 10.8±7 |
| CD68 | 3.2±2 | 5.0±2* |
| CD80 | 3.7±2 | 5.4±3* |
| CD163 | 4.1±3 | 5.5±3 |
| CD206 | 1.9±1 | 2.9±2* |
| TLR2 | 1.1±0.5 | 1.3±0.6 |
| TLR4 | 1.5±0.7 | 2.1±1* |
| LLBR1 | 1.5±2 | 2.7±3 |

To next assessed healthy dMφ phenotype compared to PE. We found that in PE, increased levels of CD68+ bMφ were present (28% vs. 43%, p<0.05, Supplementary table S3 and S4). No significant change in frequency or expression of other receptors, including CD36, CD80, CD163, TLR2, TLR4 and LILBR1, was found (Supplementary table S3 and S4).

### Preeclamptic dMφ exhibit impaired efferocytosis

We next assessed isolated dMφ efferocytosis of apoptotic trophoblasts, as a measure of the main function in dMφ in the growing placenta. In healthy placentas, 70% of bMφ efferocytosed within 90 minutes, but this was significantly impaired in PE placentas, where only 25% bMφ efferocytosed within that time (p<0.01, Figure 2A). When the amount of trophoblasts taken up was measured by MFI, a similar amount was observed between conditions, with a non-significant downward trend in PE (p>0.05, Figure 1B). In healthy placentas, 60% pMφ efferocytosed within 90 minutes, which was significantly impaired in PE placentas, where only 30% pMφ efferocytosed (p<0.05, Figure 2C). When the amount of trophoblasts taken up was measured by MFI, there was also a significant decreased by PE pMφ (4.5 vs. 2.1, p<0.05, Figure 2D). Internalisation of trophoblasts was confirmed at 10 minutes (Fig 2E), whereby small red dots of trophoblast are visible within macrophages. After 60 minutes (Fig 2F), the spread of dye was larger within macrophages, likely due to breakdown of phagolysosomes and spreading of dye within the macrophage.

**Figure 2.**
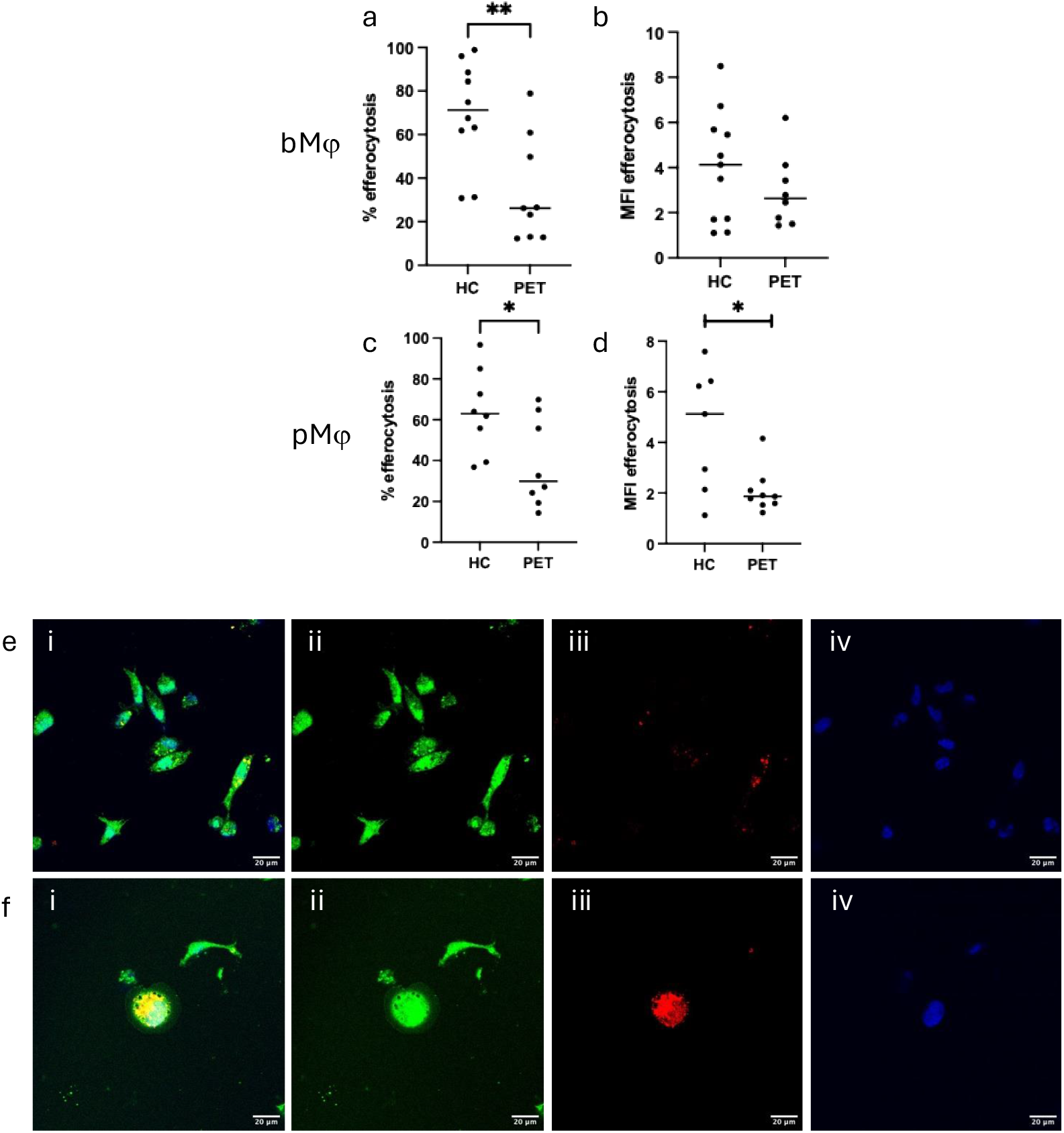
DMφ efferocytosis is impaired in PET. DMφ were incubated with apoptotic trophoblasts for 90 minutes, and uptake measured by flow cytometry. a) bMφ efferocytosis is significantly impaired in PET (HC 70±7.6% vs. PET 34±8%, p<0.05). b) The amount of trophoblasts efferocytosed measured by MFI is not significantly impaired (HC 4±0.7 vs. PET 3±0.6, p>0.05). c) pMφ efferocytosis is significantly impaired in PET (HC 64±7.3% vs. PET 39±8%, p<0.05). d) The amount of trophoblasts efferocytosed measured by MFI is significantly impaired in PET (HC 4.5±0.9 vs. PET 2±0.3, p<0.05). e) Confocal microscopy of bMφ efferocytosis at 10 minutes. ei) Merge, eii) bMφ stained with cell tracker green eiii) Bewo stained with cell tracker red eiv) DAPI f) Confocal microscopy of bMφ efferocytosis at 60 minutes. fi) Merge fii) bMφ stained with cell tracker green fiii) Bewo stained with cell tracker red, dispersed within the Mφ fiv) DAPI

### Preeclamptic dMφ secrete elevated pro-inflammatory cytokines

We next assessed cytokine release in our cells prior to (Unstimulated, US), or after 90 minutes of efferocytosis. In bMφ, CXCL-8 release was reduced by efferocytosis in HC placenta (15ng/ml vs. 8.2ng/ml, p<0.05), but not in PE placenta (25ng/ml vs. 23ng/ml, p>0.05), where release of CXCL-8 after efferocytosis was significantly increased compared to HC (p<0.05, Figure 3A). Similarly, IL-6 release was significantly elevated after efferocytosis in PE placentas, compared to HC (16pg/ml vs. 78pg/ml, p<0.05, Figure 3C). Despite this impairment to inhibit pro-inflammatory cytokine release by PE bMφ, IL-10 release was elevated in both HC (40pg/ml vs. 138pg/ml, p<0.05, Figure 3E) and PE bMφ (55pg/ml vs. 171g/ml, p<0.05) after efferocytosis.

**Figure 3.**
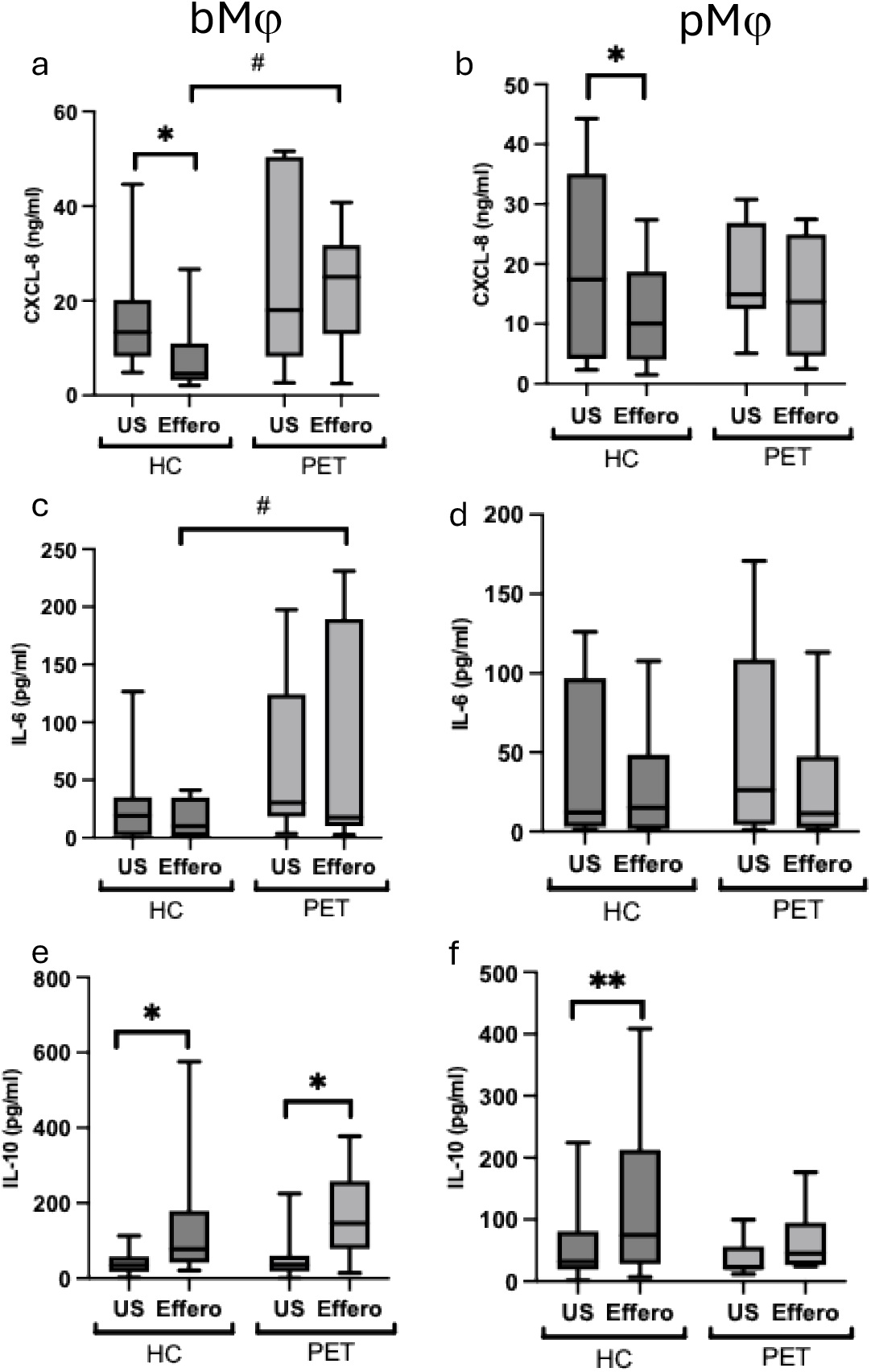
Cytokine release from DMφ after efferocytosis. DMφ were unstimulated (US) or allowed to efferocytose apoptotic trophoblasts for 90 minutes, then supernatant removed for ELISA. a) bMφ CXCL-8 release shows reduced CXCL-8 release after efferocytosis in HC (16±3ng/ml vs. 8±2ng/ml, p<0.05), but no change in PET (26±7.5ng/ml vs. 23±4.5ng/ml, p>0.05. PET bMφ release significantly increased CXCL-8 after efferocytosis then HC (p<0.05). b) pMφ CXCL-8 release shows reduced CXCL-8 release after efferocytosis in HC (20±5ng/ml vs. 11±2.6ng/ml, p<0.05), but no change in PET (18±3ng/ml vs.14±3.5ng/ml, p>0.05). c) bMφ IL-6 release shows no change in IL-6 release after efferocytosis by HC (26±9ng/ml vs.16±4ng/ml, p>0.05) or PET (71±27ng/ml vs.78±36ng/ml, p>0.05. PET bMφ release significantly increased IL-6 after efferocytosis then HC (p<0.05). d) pMφ IL-6 release shows no change across treatments or patient groups. e) bMφ IL-10 release shows increased IL-10 release after efferocytosis in HC (41±9ng/ml vs. 138±42ng/ml, p<0.05), and in PET (56±25ng/ml vs. 171±41ng/ml, p<0.05). There was no difference between patient groups. f) pMφ IL-10 release shows increased IL-10 release after efferocytosis in HC (63±25ng/ml vs. 130±42ng/ml, p<0.01), but not in PET (37±11ng/ml vs. 65±19ng/ml, p<0.05. There was no difference between patient groups.

In pMφ, a similar reduction in CXCL-8 release was observed in HC macrophages (19ng/ml vs. 11ng/ml, p<0.05, Figure 2B), which was not emulated in PE macrophages (18pg/ml vs. 15pg/ml, p>0.05, Figure 2B). IL-6 release did not change between groups (HC 30ng/ml vs. 29ng/ml, p>0.05 and PE 50pg/ml vs. 40pg/ml, p>0.05, Figure 3d). IL-10 release was elevated in HC pMφ after efferocytosis (62pg/ml vs. 130pg/ml, p<0.01, Figure 3f), but not by PE macrophages (37pg/ml vs. 65pg/ml, p>0.05, Figure 3f), indicating a failure of response.

### dMφ efferocytosis elevate ‘eat me’ receptors on the cell surface

We next assessed expression of ‘eat me’ receptors by macrophages after efferocytosis, across pooled patient groups. Our gating strategy is in Supplementary figure S1. Untreated macrophages express low levels of CD36, integrin, LLBR1, MERTK, TIM1 and TIM4, with similar levels of expression between bMφ and pMφs (Figure 4 a-f). In Mφ that had successfully efferocytosed (Effero+), macrophages elevated expression of integrin (bMφ p<0.001, pMφ p<0.001, Figure 4b), LLBR1 (bMφ p<0.001, pMφ p<0.01, Figure 4c), TIM1 (bMφ p<0.001, pMφ p<0.01, Figure 4e) and TIM4 (bMφ p<0.05, Figure 4f), with trends seen for CD36 (p>0.05, Figure 4a) and MERTK (p>0.05, Figure 4d). However, macrophages that did not efferocytose (Effero-) fail to elevate these receptors, and they remain at baseline levels. This separation in responses is similar across both dMφ locations, and suggests that elevated receptor expression primes for further efferocytosis in dMφ capable of this process.

**Figure 4.**
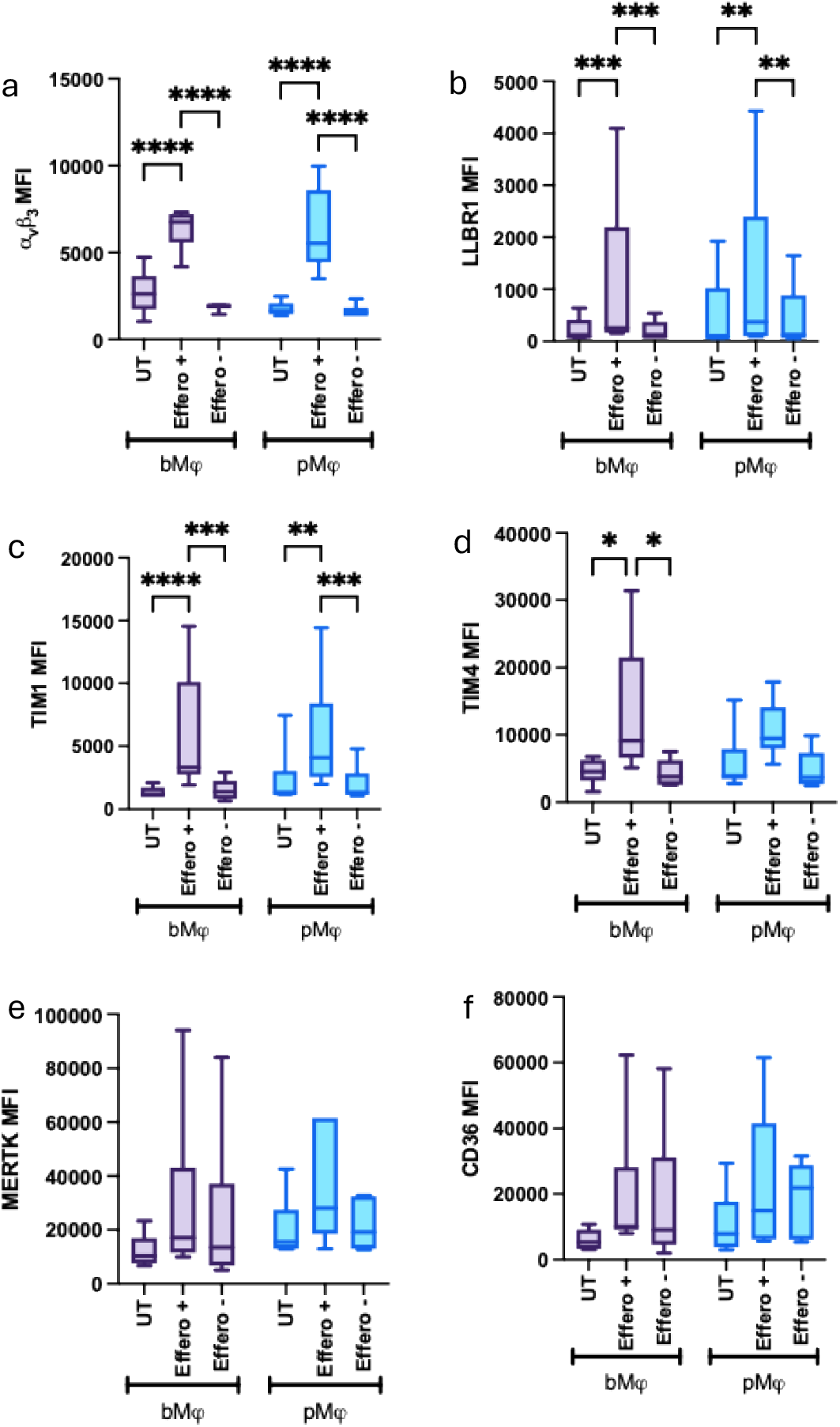
DMφ expression of efferocytosis receptors. bMφ or pMφ were left untreated (UT) or allowed to efferocytose apoptotic trophoblasts for 90 minutes, then uptake and expression of receptors measured by flow cytometry. DMφ are gated based on whether they have efferocytoise (Effero+) or not (Effero -) within the same cell population. a) Integrin α_v_β_3_ expression is elevated in Effero+ Mφ, but remains low in Effero-Mφ for both bMφ (UT 2710±522 vs. effero+ 6371±480 (p<0.001) vs effero-1871±87 (p<0.001)) and pMφ (UT 1767±162 vs effero+ 6235±977 (p<0.001) vs. effero - 1635±149 (p<0.001). b) LLBR1 expression is elevated in Effero+ Mφ, but remains low in Effero-Mφ for both bMφ (UT 215±107 vs. effero+ 991±777 (p<0.001) vs effero-200±87 (p<0.001)) and pMφ (UT 450±369 vs effero+ 1072±841 (p<0.01) vs. effero - 404±310 (p<0.01). c) TIM-1 expression is elevated in Effero+ Mφ, but remains low in Effero-Mφ for both bMφ (UT 1331±185 vs. effero+ 5796±1997 (p<0.001) vs effero-1539±341 (p<0.001)) and pMφ (UT 2315±1031 vs effero+ 5627±1871 (p<0.01) vs. effero - 1979±588 (p<0.001). d) TIM-4 expression is elevated in Effero+ Mφ, but remains low in Effero-Mφ for bMφ (UT 4551±775 vs. effero+ 13358±4050 (p<0.05) vs effero-4343±839 (p<0.05)) but not for pMφ (UT 5840±1900 vs effero+ 10695±1726 (p>0.05) vs. effero - 4851±1182 (p>0.05). e) There was no significant change in MERTK expression between treatments. f) There was no significant difference in CD36 expression between treatments.

### dMφ phagocytosis of Streptococcus agalactae

Finally, we assessed the ability of macrophages to phagocytose bacteria. We observed a similar impairment in bMφ (23% vs. 8.4%, p<0.05, Figure 5A) and pMφ (26% vs. 7.8%, p<0.05, Figure 5B) phagocytosis in PE macrophages compared to HC, although there was no change in MFI (p>0.05, Figure 5C, D). This resulted in no change in CXCL-8 and TNFα release by these macrophages (Supp figure 2), but we saw elevated release of IL-6 by PE bMφ (p<0.05), which was significantly elevated compared to HC bMφ (p<0.05, Figure 5e). No difference between pMφ macrophages was seen for IL-6 release (Figure 5F). Phagocytosis elevated IL-10 release by HC bMφ (p<0.05, Figure 5g), but not PE bMφ (p>0.05), which may indicate a protective mechanism against inflammation in the healthy decidua. No change in IL-10 release was seen for pMφ (Figure 5h).

**Figure 5.**
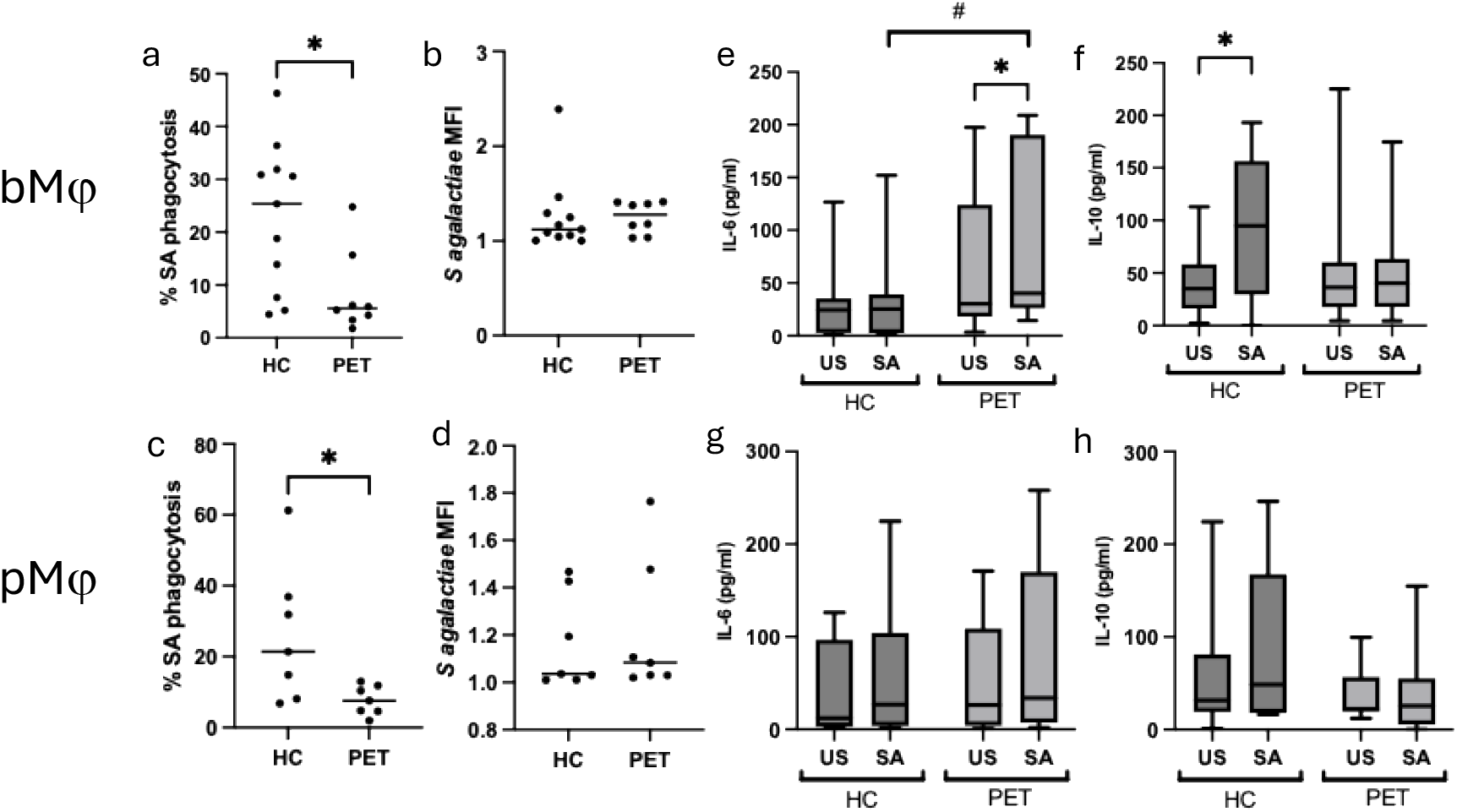
DMφ phagocytosis is impaired in PET. DMφ were incubated with heat-killed, fluorescently labelled SA for 4 hours, and uptake measured by flow cytometry. a) bMφ phagocytosis is significantly impaired in PET (HC 23±4% vs. PET 8.4±3%, p<0.05). b) The amount of SA phagocytosed measured by MFI is not significantly impaired (HC 1.3±0.1 vs. PET 1.3±0.1, p>0.05). c) pMφ phagocytosis is significantly impaired in PET (HC 26±7% vs. PET 8±1.6%, p<0.05). d) The amount of SA phagocytosed measured by MFI is not significantly impaired (HC 1.2±0.1 vs. PET 1.2±0.1, p>0.05). Cytokine release by DMφ after phagocytosis. e) bMφ release elevated IL-6 after phagocytosis in PET (71±27pg/ml vs. 97±33pg/ml, p<0.05), but no change in HC (28±10ng/ml vs. 33±12ng/ml, p>0.05). PET DMφ release significantly increased IL-6 after phagocytosis then HC (p<0.05). f) bMφ release elevated IL-6 after phagocytosis in HC (41±31pg/ml vs. 93±67pg/ml, p<0.05), but no change in PET (56±71ng/ml vs. 53±53ng/ml, p>0.05). g) pMφ show no change in the release of IL-6 after phagocytosis (p>0.05). h) pMφ show no change in the release of IL-10 after phagocytosis (p>0.05).

## Discussion

Efferocytosis at the maternal-fetal interface is a vital process to prevent the accumulation of necrotic cells that can promote inflammation, and the development of pregnancy complications. Here we demonstrate for the first time, that dMφ from both the decidua basalis, and decidua parietalis are capable of efferocytosing apoptotic trophoblasts, which reduces the release of pro-inflammatory CXCL-8 and IL-6, and induces release of anti-inflammatory IL-10, thus maintaining an anti-inflammatory, tolerogenic environment in the decidua. We also show that in patients with early-onset PE, efferocytosis of trophoblasts is impaired, associated with elevated pro-inflammatory CXCL-8 and IL-6 release. These findings are the first to implicate impaired dMφ function in the pathogenesis of PE and suggest that macrophage dysfunction can contribute to the development of pregnancy complications.

Apoptosis, or programmed cell death is a process that is initiated due to stimuli that signal for the cell to die, and differs from necrosis which is non-regulated, and results in the cell dying and releasing intracellular contents which promote inflammation [2]. Levels of apoptotic cells in the placenta increase from first to third trimester of normal pregnancy [23], and require removal by resident maternal immune cells.

In the first trimester, dMφ and uNK cells are found within the spiral arteries where they disrupt vascular smooth muscle cells (VSMC) through the release of MMP7 and 9, which may induce apoptosis of VSMC to allow trophoblast invasion of the artery and remodelling to occur successfully [24]. *Ex vivo*, dMφ have been shown to efferocytose apoptotic VSMC after 72 hours of co-culture, alongside the release of pro-angiogenic factors which promote spiral artery remodelling [25]. We hypothesis, therefore, that dMφ not only control the microenvironment through release of pro-angiogenic factors, but also allow remodelling to occur without excessive inflammation through the removal of apoptotic cells generated during this process.

As pregnancy progresses, dMφ remain anti-inflammatory and likely maintain tolerance through regular removal of apoptotic cells generated through normal cellular turnover. Our data supports this theory, indicating that in healthy pregnancy, efferocytosis of trophoblasts promotes the release of IL-10, which plays a range of roles in the placenta including promoting HLA-G expression by trophoblasts and limiting MMP-9 secretion and invasiveness of extravillous trophoblasts [26, 27], alongside anti-inflammatory effects. While this observation was more prominent in bMφ, we saw similar effects in the pMφ which are primed as more pro-inflammatory antigen presenting associated roles [11] but clearly display plasticity when exposed to apoptotic stimuli.

In preeclampsia, the placenta is in a heightened inflammatory state due to impaired spiral artery remodelling and shallow trophoblast invasion [28]. Elevated levels of apoptosis in PE have been shown by IHC and protein expression of Fas and caspase markers, which are high in syncitiotrophoblasts, cytotrophoblasts and extravillous trophoblasts [29, 30]. Our study shows impaired ability of dMφ from early onset PE placentas to efferocytosis apoptotic trophoblasts, which may have profound implications in the pathogenesis of this disease. Firstly, while our data stem from third trimester samples, if this process begins early in pregnancy, the impaired efferocytosis of trophoblasts and VSMC generated during spiral artery remodelling and trophoblast invasion may die by necrosis and generate inflammation which impairs the correct formation of the placenta – promoting placental dysfunction as pregnancy progresses. Secondly, whether this dMφ dysfunction occurs early or late in gestation, the impaired removal of apoptotic cells will promote the release of DAMPs, ATP and other cellular contents which drive the increase in inflammatory and oxidative stress mediators seen the placenta of preeclampsia [31]. Secondary necrosis subsequent to impaired efferocytosis releases cell free DNA and G-actin into the extracellular space, which can enter the maternal blood and trigger damage to the maternal endothelium, contributing to the onset of hypertension [32]. Overall, impaired efferocytosis and elevated CXCL-8 and IL-6 release by dMφ likely drives inflammation and contributes to the pathogenesis of this disease.

We attempted to understand the process of efferocytosis by dMφ, as the first step to understanding how dMφ function, and how they might be implicated in preeclampsia. We assessed the expression of ‘eat me’ receptors on the dM surface and identified significant differences between macrophages that efferocytose, and those that do not. In untreated dMφ, expression of integrin, TIM1, TIM4, MERTK, CD36 and the trophoblast receptor LLBR1 was low but similar across macrophage location. However, on exposure to apoptotic trophoblasts, efferocytic macrophages elevated their expression of these receptors. This is likely due to priming of the macrophage, as once one apoptotic cells has bound to receptors, the cell mobilises stored of receptors to elevate its ability to efferocytose further [33]. The consequence of receptor elevation is likely to be further and prolonged efferocytosis, which is beneficial in the placenta to prevent inflammation caused by dying cells [34].

However, we noted that non-eaters fail to mobile receptors, and levels remain the same as unstimulated macrophages. This failure to efferocytose may be indicative of a number of consequences. It is possible that there are multiple functions of dMφ, and that a portion of the macrophage pool do not efferocytose, but maintain another unknown function. Another theory is that some macrophages are exhausted, and therefore lose the capacity to take up further cells [35] [36]. This may be the case in these *ex vivo* macrophages, which will have been exposed to efferocytosis in the placenta, and therefore may be non-functional *ex vivo*. This may have implications in our study, wherby PE macrophages fail to efferocytose, which could be due to an innate mechanism that requires further understanding, or may be due to exhaustion *in vivo*, which cannot be recapitulated when isolated *ex vivo*. Further assessment of this phenomenon is required to fully understand both the mechanism of impaired efferocytosis in PE dMφ, and the process of efferocytosis in the placenta.

Finally, we assessed the ability of dMφ to phagocytose GBS, the main infection causing bacteria of the placenta, and observed a similar defect in uptake by PE macrophages in both the bMφ and pMφ. This likely indicates a shared mechanistic defect in uptake by PE dMφ, which again requires further evaluation. However, it does suggest that patients with PE, or other causes of placental damage, may be more likely to develop a GBS infection during pregnancy, which would further increase the risk to the infant.

We recognise some limitations to our study. While our patient numbers are small, we were able to detect statistically significant differences in dMφ function, but our analysis is limited to functional outputs. Further linked assessment of generalised placental inflammation would strengthen our understanding of dMφ in this context. Our samples display a gestational mismatch of 5 weeks, with the PE samples delivered at early gestation, and with smaller babies. This could have some implication for our results, however we must recognise that this represents the significant impact of PE on pregnancy outcomes, and that gestational matching is not possible with healthy pregnancies.

In conclusion, we show that impaired dMφ efferocytosis of trophoblasts in early onset PE is associated with pro-inflammatory cytokine release which may drive inflammation in the placenta. Whether this phenomenon is implicated in impaired spiral artery remodelling and trophoblast invasion in early pregnancy or is limited to later in pregnancy where it contributes to increased inflammation, oxidative stress and endothelial damage remains to be determined. The origin of altered dMφ function also remains unexplored, with potential pre-pregnancy endometrial changes possible that prelude to the onset of PE or other pregnancy complications. Further research into this phenomenon is warranted, as targeting macrophage function in the decidua may provide a useful therapeutic strategy to prevent or limit the severity of preeclampsia.

## Data availability statement

The data that support this study are not openly available due to reasons of sensitivity and are available from the corresponding author upon reasonable request.

## Acknowledgements

Authors would like to thank the University of Manchester Flow Cytometry Core Facility, and the University of Birmingham Flow Cytometry Platform for assistance with flow cytometry. We would also like to thank the University of Manchester Bioimaging facility for assistance with confocal microscopy. Finally, we would like to thank the research practitioners and midwives at Birmingham Women’s Hospital, and St Marys Hospital Birmingham for assistance with patient recruitment and sample collection.

