## Supplementary figure for "Impaired trophoblast efferocytosis by decidual macrophages in early-onset preeclampsia"

#### Supplementary table S1

Antibodies for DM phenotyping. All antibodies were from Biolegend

| Marker | Fluorophore | Order code | Volume µL |
|---|---|---|---|
| CD14 | BV421 | 367144 | 2.5 |
| CD10 | BV510 | 312220 | 2.5 |
| CD206 | BV605 | 321140 | 2.5 |
| CD64 | BV650 | 305054 | 2.5 |
| CD80 | BV711 | 305236 | 2.5 |
| CD68 | BV785 | 333826 | 2.5 |
| LLBR1/CD85j | FITC | 333730 | 2.5 |
| TLR4 | PE | 312806 | 2.5 |
| CD163 | PE Cy5 | 333644 | 2.5 |
| TLR2 | PE Cy7 | 309708 | 2.5 |
| HLA-DR | APC | 980406 | 2.5 |
| CD36 | AF700 | 336236 | 2.5 |

#### Supplementary table S2

Antibodies for efferocytosis. All antibodies were from Biolegend, and cell tracker green CMFDA was from thermofischer.

| Marker | Fluorophore | Order code | Volume |
|---|---|---|---|
| CD14 | BV421 | 367144 | 2.5 |
| CD10 | BV510 | 312220 | 2.5 |
| MERTK | BV711 | 367620 | 2.5 |
| BeWo | Cell tracker green | C2925 | 1 |
| Int a5b3 | PE | 304406 | 2.5 |
| LILRB1 | PE-dazzle594 | 333716 | 2.5 |
| TIM1 | PerCP Cy5.5 | 353912 | 2.5 |
| TIM4 | PE Cy7 | 354006 | 2.5 |
| HLA-DR | APC | 980406 | 2.5 |
| CD36 | AF700 | 336236 | 2.5 |

### Supplementary table S3

% receptor expression in healthy vs PET macrophages

| | Basalis | | Parietalis | |
|---|---|---|---|---|
| | HC | PET | HC | PET |
| CD36 | 46±21 | 41±14 | 38±21 | 38±17 |
| CD68 | 28±15 | 43±8* | 40±23 | 57±31 |
| CD80 | 37±12 | 40±13 | 51±22 | 62±20 |
| CD163 | 79±25 | 75±30 | 81±27 | 83±20 |
| CD206 | 40±21 | 41±30 | 53±24 | 61±19 |
| TLR2 | 41±23 | 46±19 | 47±18 | 59±14 |
| TLR4 | 87±14 | 77±27 | 86±23 | 86±18 |
| LLBR1 | 22±19 | 25±9 | 32±28 | 51±28 |

### Supplementary table S4

**M**FI of receptor expression in healthy vs PET macrophages

| | Basalis | | Parietalis | |
|---|---|---|---|---|
| | HC | PET | HC | PET |
| CD36 | 10.6±6 | 9.8±6 | 10.8±7 | 18±14 |
| CD68 | 3.2±2 | 4.2±3 | 5.0±2 | 11±9 |
| CD80 | 3.7±2 | 4.5±4 | 5.4±3 | 9±7 |
| CD163 | 4.1±3 | 5±2 | 5.5±3 | 10±7 |
| CD206 | 1.9±1 | 2.3±1 | 2.9±2 | 3.8±2 |
| TLR2 | 1.1±0.5 | 1.9±1 | 1.3±0.6 | 3±2.6 |
| TLR4 | 1.5±0.7 | 2.6±2 | 2.1±1 | 3.5±2 |
| LLBR1 | 1.5±2 | 1.5±1 | 2.7±3 | 4±5 |

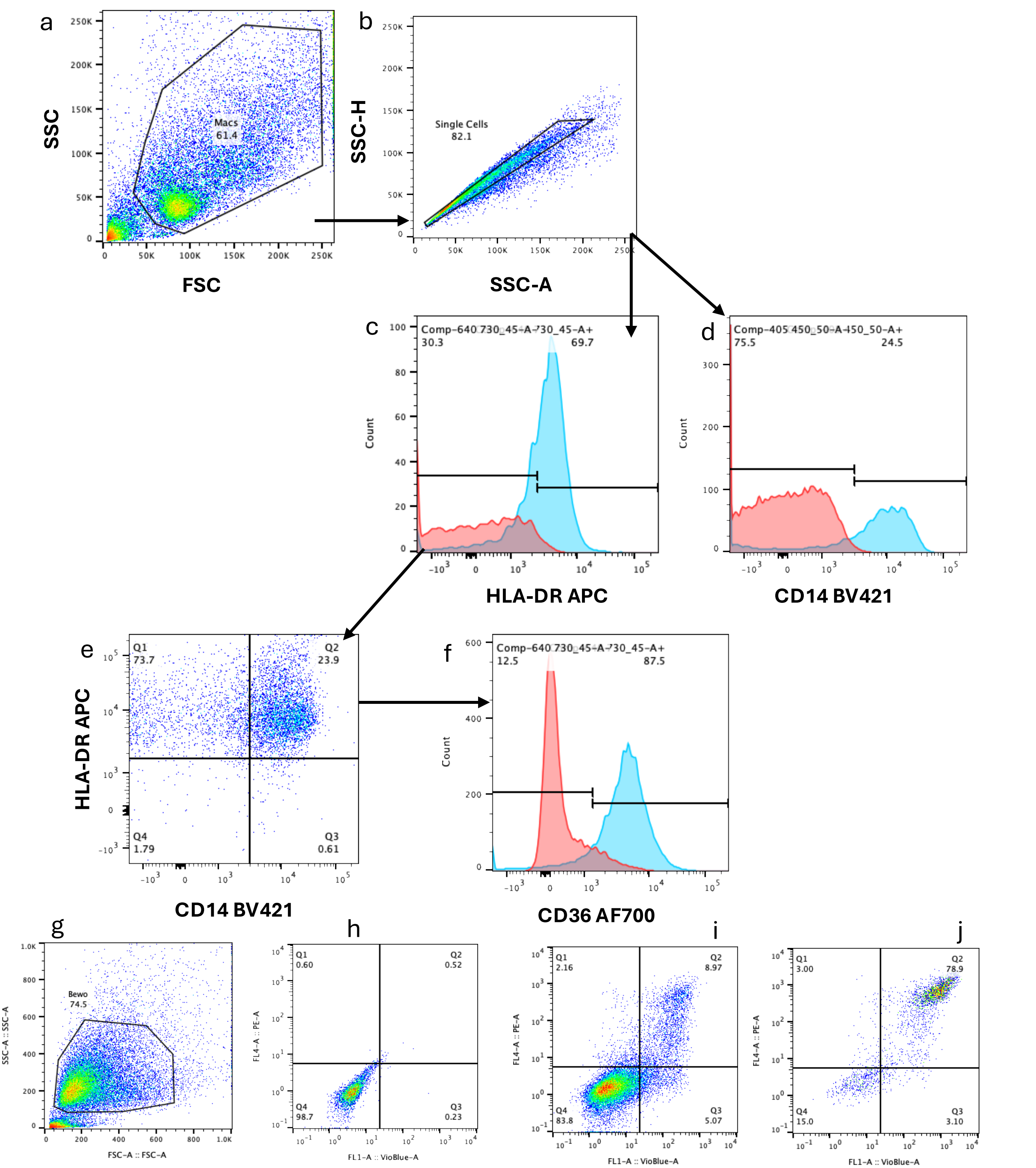


**Supplementary figure 1 – Gating strategy**

a) Isolated DMφ were gated based on FSC vs SSC plot. b) Single cells were gates based on SSC-A vs SSC-H. DMφ were identified based on HLA-DR+ CD14+ - c) shows HLA-DR+ gating (blue) vs. FMO control (red). d) shows CD14+ gating (blue) vs. FMO control (red). e) Resulting CD14+ HLA-DR+ DMφ identified in Q2 gate. f) Example phenotyping gating for CD36 based on positive sample (blue) vs. FMO control (red).

Bewo Apoptosis – g) Bewo cells were gated based on FSC vs SSC plot. h) An unlabelled sample was gated as negative for Annexin V-BV421 vs. Propidium idodide (PE channel) expression. i) A live sample of Bewo cells shows 84% live (Q4 gate), 5% early apoptotic (Q3 gate), 9% late apoptotic (Q2 gate, and 2% dead (Q1 gate). j) UV-treated Bewo show 80% late apoptotic (Q2 gate).

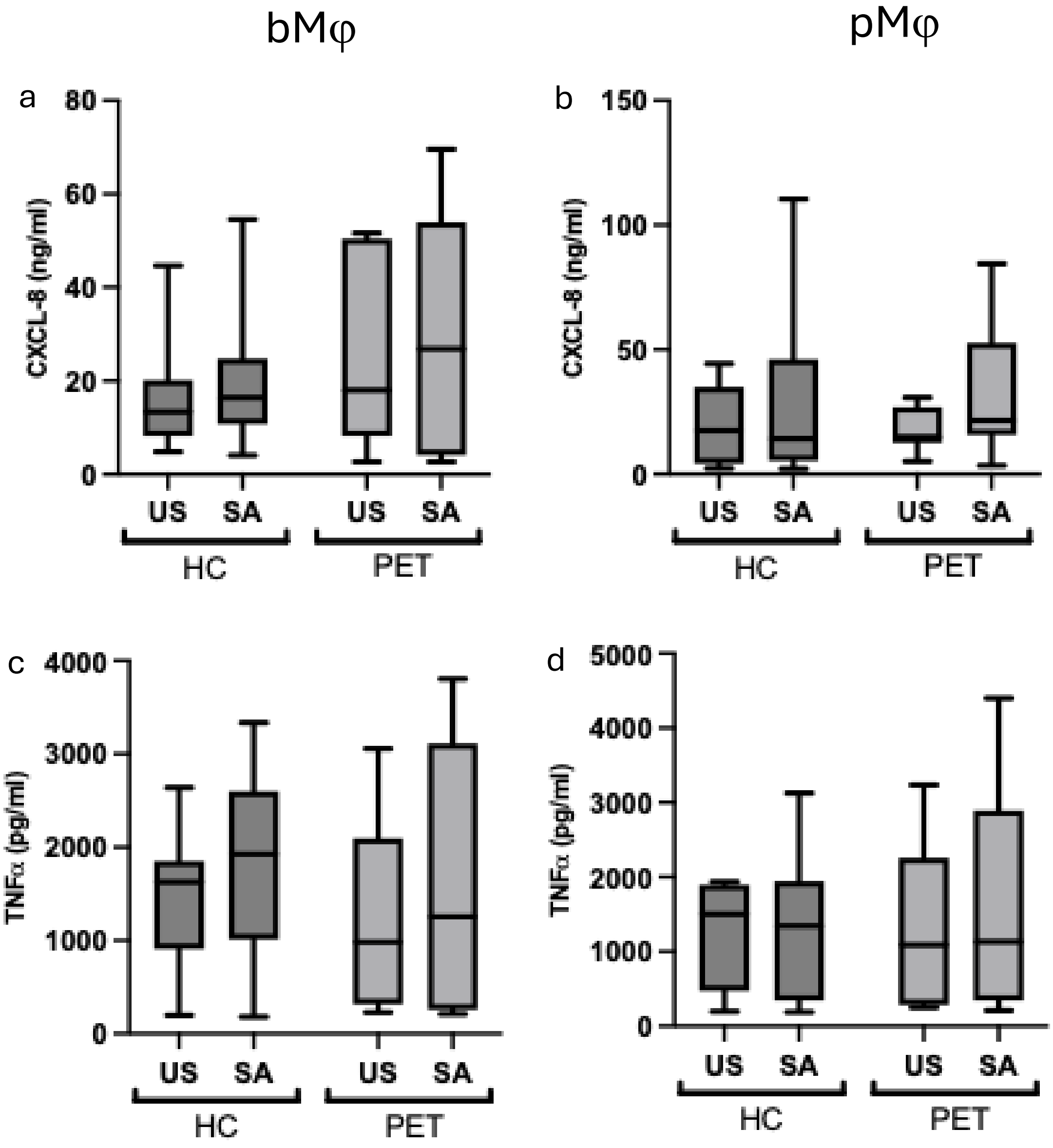


**Supplementary figure 2 - Cytokine release from DMφ after phagocytosis**

DMφ were incubated with heat-killed, fluorescently labelled SA for 4 hours, and uptake measured by flow cytometry. There was no change in the release of CXCL-8 from both bMφ (a) or pMφ (b), or the release of TNFα from both bMφ (c) or pMφ (d)
